# PixR, a novel activator of conjugative transfer of IncX4 resistance plasmids, mitigates the fitness cost of *mcr-1* carriage in *Escherichia coli*

**DOI:** 10.1101/2021.08.04.455183

**Authors:** Lingxian Yi, Romain Durand, Frédéric Grenier, Jun Yang, Kaiyang Yu, Vincent Burrus, Jian-Hua Liu

## Abstract

The emergence of the plasmid-borne colistin resistance gene *mcr-1* threats public health. IncX4-type plasmids are one of the most epidemiologically successful vehicles for spreading *mcr-1* worldwide. Since MCR-1 is known for imposing a fitness cost to its host bacterium, the successful spread of *mcr-1*-bearing plasmids might be linked to high conjugation frequency, which would enhance the maintenance of the plasmid in the host without antibiotic selection. However, the mechanism of IncX4 plasmids conjugation remains unclear. In this study, we used high-density transposon mutagenesis to identify factors required for IncX4 plasmid transfer and 18 genes were identified, including five with annotations unrelated to conjugation. The Cappable-seq and RNA-seq analysis confirmed that a novel transcriptional regulator gene, *pixR*, directly regulates the transfer of IncX4 plasmids by binding the promoter of 13 essential transfer genes to increase their transcription. Plasmid invasion and co-culture competition assays revealed that *pixR* is essential for the spread and persistence of *mcr-1*-bearing IncX4 plasmids in bacterial populations, and effective conjugation is crucial for alleviating the fitness cost exerted by *mcr-1* carriage. The existence of the IncX4-specific *pixR* gene increases plasmid transmissibility while promoting the invasion and persistence of *mcr-1*-bearing plasmids in bacterial populations, which helps explain their global prevalence.

**IMPORTANCE:** The spread of clinical important antibiotic resistance genes is frequently related to some epidemic plasmids. However, the underlying molecular mechanisms contributing to the successful spread of these epidemic plasmids remains unclear. The significant of our research indicated that efficient conjugation could promote the invasion and persistence of plasmids within a bacterial population, resulting in the successful dissemination of epidemic plasmids in nature. Our data also highlight the importance of developing plasmid conjugation inhibitors to solve the antibiotic resistance crisis.

## INTRODUCTION

Plasmids play a significant role in the evolution of antibiotic resistance (1). Plasmid-borne antibiotic resistance genes can efficiently spread within bacterial communities across diverse species, driving antibiotic resistance evolution and threatening our ability to treat bacterial infections by restricting treatment options (2). The rise of carbapenem-resistant *Enterobacteriaceae* (CRE) and extended-spectrum beta-lactamase (ESBL)-producing *Enterobacteriaceae* is the most alarming example. The World Health Organization (WHO) listed these microorganisms as critical pathogens for which the development of new treatments is a high priority due to the global spread of plasmid-borne carbapenem resistance and ESBL genes (3). Colistin is one of a few effective options to treat infections caused by CRE. However, the emergence and dissemination of the plasmid-borne colistin resistance gene *mcr-1* limit the efficacy of colistin, causing worldwide concern (4).

Previous work revealed IncX4 plasmids are the second most prevalent epidemic vectors carrying *mcr-1* after IncI2 plasmids (5–7). As of May 2021, IncX4 plasmids carrying *mcr-1* have been identified in *Escherichia coli, Salmonella enterica* (serovars Typhimurium, Paratyphi B, Java, Anatum, and Schwartzengrund), *Klebsiella pneumoniae*, and *Enterobacter cloacae* isolated from humans, food products, livestock, wildlife, and environmental samples across more than 41 countries and regions (Figure S1A, Table S1) (8–11). Besides *mcr-1*, IncX4 plasmids are also associated with the transmission of the 23S rRNA methyltransferase gene *cfr*, and the extended spectrum β-lactamase and carbapenemase genes *bla*_TEM_, *bla*_CTX-M_, *bla*_OXA_, and *bla*_NDM_ (12, 13).

Bacterial immunity systems, such as CRISPR-Cas or restriction-modification (RM) systems, can restrict the entry and establishment of antibiotic resistance plasmids that are recognized as exogenous DNA. These plasmids also inflict a fitness cost on the host, leading to their loss in the absence of selective pressure (14–16). Paradoxically, in natural settings, epidemic antibiotic resistance plasmids can persist in bacterial populations over the long term without antibiotic selection. The latest theory explaining this “plasmid-paradox” examines three main features: the rate of plasmid loss during replication, the rate of plasmid acquisition, and the plasmid fitness adaptation to the host (1, 17, 18). Conjugation plays an essential role in the persistence of a plasmid in a bacterial population (19–21). Previous reports have shown that IncX4 plasmids transfer at a high frequency, ~10^-2^-10^-4^ (22, 23). Although the rapid transmission of *mcr-1*-bearing IncX4 plasmids sparked intense concerns, most of the research conducted to date has only focused on their epidemiology. Hence, the mechanisms that have enabled IncX4 plasmids to become successful vectors for the global spread of *mcr-1*, to adapt to hosts such as *E. coli*, and to maintain antibiotic resistance genes in natural bacterial communities are unclear. The conjugative transfer genes (*taxB-pilX-taxCA*) of IncX4 plasmids were predicted based on sequence similarity with the IncX family reference plasmid R6K. R6K encodes a P-type (*virB*) type IV secretion system (T4SS), PilX1-PilX11, involved in pilus synthesis and assembly. It also encodes the relaxase TaxC, the auxiliary factor TaxA, and the coupling protein TaxB, which are involved in DNA processing during conjugation (GenBank accession no. AJ006342) (24). However, the genes carried by IncX4 plasmids share limited sequence identity (<60%) with those of R6K (23). Hence, the determinants of IncX4 plasmid transfer remain largely unknown, and the regulation of their transfer is poorly understood. Plasmid transfer inflicts a fitness cost due to the high ATP demand to synthesize the mating channel and energize the translocation of plasmid DNA into the recipient cell (25). To minimize this burden, the expression of the conjugative transfer genes is usually tightly controlled by plasmid- and host-encoded factors (16). The factors involved in the regulation of IncHI, IncA, IncC, IncP, and F-like plasmids are now well characterized (26–29). In contrast, little is known about the control of IncX conjugation.

Here, we used a systematic approach to identify the genes required for IncX4 plasmid transfer and characterized an IncX4-specific transcriptional activator, PixR, that controls plasmid conjugation. Our results indicate that efficient conjugation alleviates the fitness cost of low-copy *mcr-1* carriage and promotes the persistence and invasion of *mcr-1*-bearing plasmids within a bacterial population, which helps explain the successful dissemination of *mcr-1*-bearing plasmids in nature.

## METHODS

### Bacterial strains, plasmids and culture conditions, plasmids and strains constructions and bacterial conjugation experiments

The strains and plasmids used in this study are listed in Table S2. The primers used in this study are listed in Table S3. The detail of culture conditions, plasmids and strains constructions and bacterial conjugation experiments was described in Text S1.

### Comparative analyses

A BLASTn search (https://blast.ncbi.nlm.nih.gov/Blast.cgi) for all complete IncX4 plasmid sequences in the NCBI Genbank database was conducted using the *repA* sequences of IncX4 plasmids extracted from the PlasmidFinder database (https://bitbucket.org/genomicepidemiology/plasmidfinder_db/src/master/). A total of 247 IncX4 plasmid sequences were collected (Table S1). BLASTp and Pfam (http://pfam.xfam.org) were used to analyze the PixR protein. Protein sequences were aligned using MUSCLE 3.8.31 (30). Gene organization diagrams were generated with Easyfig 2.2.2.

### Competition experiments in *vitro*

The strains BW25113/pHNSHP23, BW25113/pHNSHP23Δ*pixR::cat* and BW25113/pHNSHP23Δ*pixR*Δ*mcr-1*::cat were used to compete against BW25113, BW25113/pHNSHP23Δ*pixR*Δ*mcr-1*::cat was used to compete against BW25113/pHNSHP23Δ*mcr-1*::cat and the strain BW25113/pHNSHP23Δ*mcr-1*::*cat* was used to compete against BW25113/pHNSHP23. All competition assays were carried out in biological triplicates. The overnight cultures of the two competitors were mixed in equal volumes. Cultures were grown for 5 days with a 1:1 000 dilution into fresh LB broth every 24 h. At each 24h timepoint, cultures were serially diluted and plated on LB agar plates containing colistin or chloramphenicol. Considering BW25113/pHNSHP23Δ*pixR*Δ*mcr-1*::*cat* and BW25113/pHNSHP23Δ*mcr-1*::*cat* could both be present on Cm plates, we distinguished these two strains by colony PCR with primers pixRc_F/R. The following formula was used to calculate the relative fitness: RF = ln(N_f,s1_/N_i,s1_)/ln(N_f,s2_/N_i,s2_). RF is the related fitness of S1 strain compared to S2 strain. N_f,sx_ and N_i,sx_ correspond to the CFU counts of strain Sx at the appropriate timepoint and at the initial timepoint, respectively.

### Plasmid invasion assays

Plasmid invasion assays were performed based on the protocol described previously with some modifications (21). Briefly, overnight cultures of the recipient strain BW25113 were diluted 1:100 in LB medium and mixed with a 1:10 000 dilution of the overnight cultures of the donor strain either BW25113/pHNSHP23, BW25113/pHNSHP45 BW25113/pHNSHP23Δ*pixR::cat*, or BW25113/pHNSHP23Δ*pixR*Δ*mcr-1::cat*. Cultures were grown in 50 mL tubes containing 2 mL LB at 37 °C with slow rolling (80 r.p.m). Every 24 h, cultures were diluted 1:100 into fresh LB broth. Viable counts were gathered at 24, 48, 72, 96,120, 144 and 168 h by plating serially diluted cultures on non-selective, Cm- or Cl-containing LB agar. CFU counts for each type of plates were used to estimate the relative quantity of cells from the donor and recipient strains. For BW25113/pHNSHP23Δ*pixR*Δ*mcr-1::cat* and BW25113/pHNSHP23Δ*pixR::cat*, we used the number of CFUs counted on Cm plates. For BW25113/pHNSHP23, we used the number of CFUs counted on Cl plates. Finally, for BW25113 we subtracted the CFUs counted on Cl plates or Cm plates from the ones counted on non-selective plates. Competitive experiments were performed in a similar fashion by co-inoculating three strains: either BW25113, BW25113/pHNSHP23 and BW25113/pHNSHP23Δ*pixR::cat*, or BW25113, BW25113/pHNSHP23, and BW25113/pHNSHP23Δ*pixR*Δ*mcr-1::cat* or BW25113, BW25113/pHNSHP23 and BW25113/pHNSHP45. Counts for BW25113/pHNSHP23 were obtained by subtracting CFUs counted on Cm plates from CFUs counted on Cl plates. To distinguish between BW25113/pHNSHP23 and BW25113/pHNSHP45 (which would both grow on CI plates), colonies were tested by PCR with primers IncI2-F/R and IncX4-F/R (Table S3).

### Cappable-seq, RNA-seq assays and High-density transposon mutagenesis (HDTM) assays

A thorough description of Cappable-seq, RNA-seq and HDTM assays procedures was given in Text S1.

### Reverse transcription and qPCR

Overnight cultures of BW25113/pHNSHP23, BW25113/pHNSHP23Δ*pixR*, BW25113/pHNSHP23Δ*pixR* + pBAD30 and BW25113/ pHNSHP23Δ*pixR* + pBAD-*pixR* were diluted to an OD_600_ of 0.05 in 2 mL LB containing the appropriate antibiotics. The cells were grown at 37°C until the OD_600_ reached 0.8 with 0.2% arabinose to induce *P_BAD_*. Total RNA was extracted by using the Hipure Bacterial RNA Kit (Magen, China). Reverse transcription was carried out with 1 μg RNA using TB Green Premix Ex Taq™ II (Takara, Japan). Targets of around 110 bp were amplified with primers qtaxB-F/R, qpilX11-F/R, qpilX3-4-F/R, and qtrbM -F/R by qPCR, while 16S expression level was used as internal control with primers 16S-F/16S-R. Relative expression was estimated by the 2^-ΔΔCt^ method. The experiment was performed with three biological replicates.

A reverse transcription experiment was performed using a primer located at the 3’ end of *trbM* and total RNA extracted from BW25113 cells bearing pHNSHP23. A PCR amplification using the reverse cDNA with primers hyp9-pilX1R/F and trbM-pilX11-R/F was performed to confirm that *cds9, trbM*, and *taxB* are indeed part of the *pilX* operon.

### Protein purification, Electrophoretic Mobility Shift Assay (EMSA) and β-galactosidase assay

The detail of protein purification, EMSA and β-galactosidase assays was described in Text S1.

### Statistical analyses and figures

Prism 9 (GraphPad Software) was used to generate graphics and perform statistical analyses. Figures were prepared with Inkscape 1.0.1 (https://inkscape.org/) and BioRender (https://biorender.com/).

### Data availability

Complete data from aligned reads for HDTM, Cappable-seq and RNA-seq experiments can be visualized using the UCSC genome browser at http://ucscbrowser.genap.ca/cgi-bin/hgHubConnect?hgHub_do_redirect=on&hgHubConnect.remakeTrackHub=on&hgHub_do_firstDb=1&hubUrl=https://datahub-102-cw2.p.genap.ca/pHNSHP23/hub.txt

## RESULTS

### Identification of the gens involved in the transfer of IncX4 plasmids

pHNSHP23 is an *mcr-1*-bearing IncX4 plasmid found in an *E. coli* strain of pig origin (6). The nucleotide sequence of pHNSHP23 was compared to a total of 246 IncX4 plasmids from diverse geographical origins found in the Genbank database using Blastn. We found that the IncX4 plasmids have a common structure with the predicted conjugative transfer locus sharing an average of 96% identity over 15 kb. Hence, we used pHNSHP23 as the prototype IncX4 plasmid for identifying all the transfer genes of IncX4 plasmids. High-density transposon mutagenesis (HDTM) was used to construct Tn*5* insertion mutant libraries of pHNSHP23 in *E. coli* BW25113. In the initial mutant library (input library), Tn*5* insertions distributed evenly across pHNSHP23, except in the gene *pir* required for IncX4 replication, the recently identified antitoxin-encoding gene *tsxB* involved in IncX4 plasmid maintenance, and the selection marker *mcr-1* (Figure 1). The input library was then used in a mating assay on agar with *E. coli* MG1655 as the recipient strain. The resulting transconjugants were used for a second round of transfer on agar (output library). Mapping of the Tn*5* insertions in the output library revealed eighteen genes identified as essential factors for conjugation, including the 10 *pilX* genes, the relaxase-encoding gene *taxC*, and DNA processing genes *taxA* and *taxB* (Figures 1 and 2A). In addition, five genes previously regarded as unrelated to conjugation were identified as potentially essential for IncX4 plasmid transfer, including the recently reported toxin-encoding gene *tsxA*, and four genes of unknown function: *cds4, cds9, cds10* and *cds16* (Figure 1 and 2A). To confirm the impact of these four genes on conjugation, we constructed individual null mutants and tested their transfer by conjugation in broth. We observed that the transfer of pHNSHP23Δ*cds9* was abolished. Compared to wild-type pHNSHP23, the transfer frequency of the Δ*cds4* and Δ*cds16* mutants was reduced ~4 fold, which was not statistically significant, whereas the transfer frequency of pHNSHP23Δ*cds10* was reduced ~100 fold (Figure 2B). These results confirm the role of the two uncharacterized genes *cds9* and *cds10* in the conjugative transfer of IncX4 plasmids.

**Figure 1.**
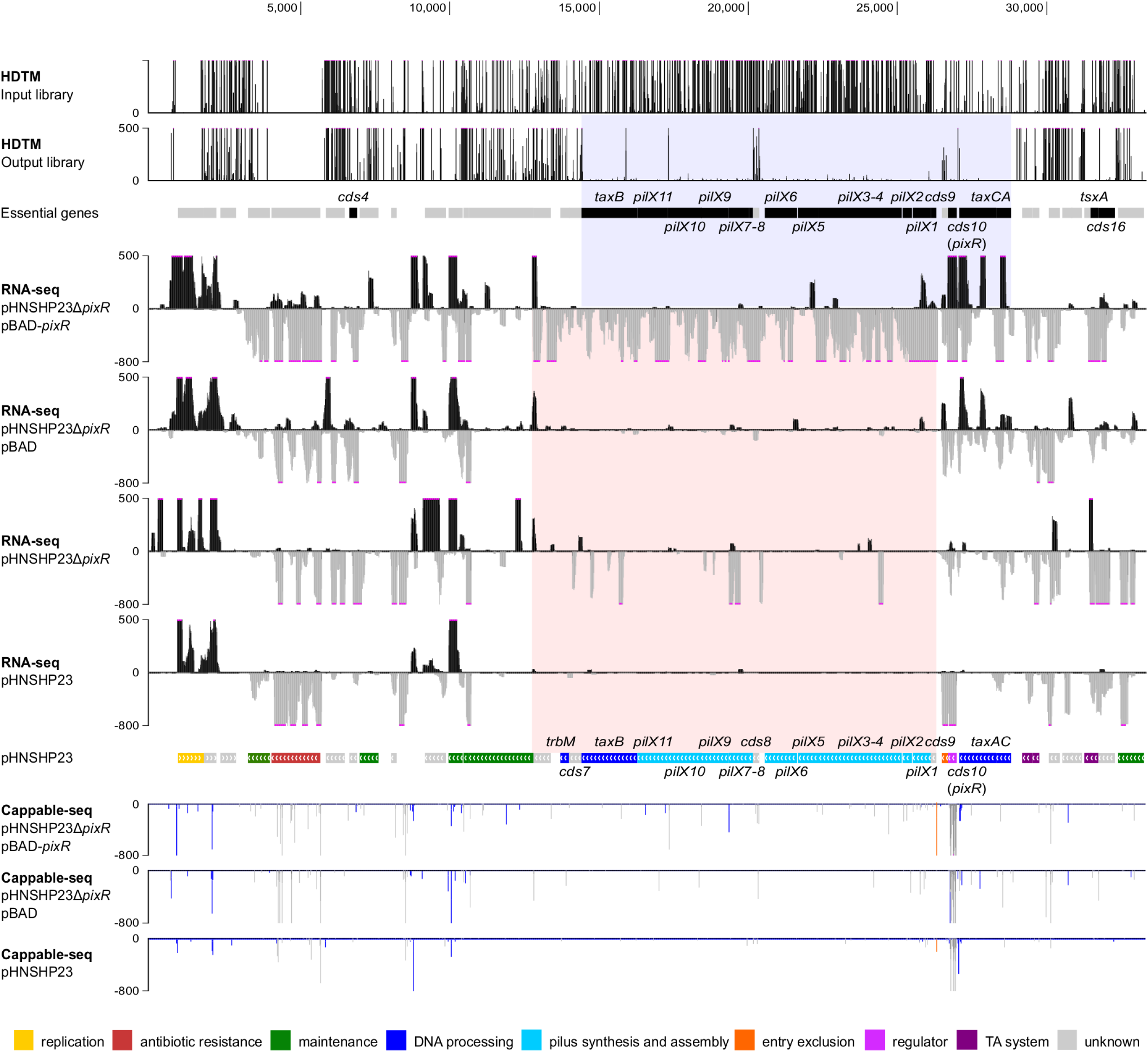
In-depth analysis of the PixR regulon. The first two tracks represent the Tn*5* insertions sites in the mutant libraries before and after two consecutive mattings, respectively. Essential genes appear in black in the ‘essential genes’ track. The region highlighted in blue encompasses the ones belonging to the transfer region. The following four tracks represent the RNA-seq read densities of *E. coli* BW25113/pHNSHP23 and BW25113/pHNSHP23Δ*pixR* carrying pBAD or pBAD-*pixR* with 0.2% arabinose induction. For each of these tracks, densities with a positive value correspond to the positive DNA strand, whereas densities with a negative value correspond to the negative DNA strand. The region highlighted in red encompasses all genes in the transfer region showing a drastic change in expression. The eighth track is the genetic map of pHNSHP23. Genes are color-coded according to their predicted function as indicated in the legend. The last three tracks represent the Cappable-seq results. Strains are detailed on the left. Blue and gray bars indicate the Cappable-seq density on the positive and negative DNA strands, respectively. The orange bar indicates the TSS of the *pilX* operon. For all tracks, pink dots indicate the signal is beyond the fixed Y-axis maximal or minimal value.

**Figure 2.**
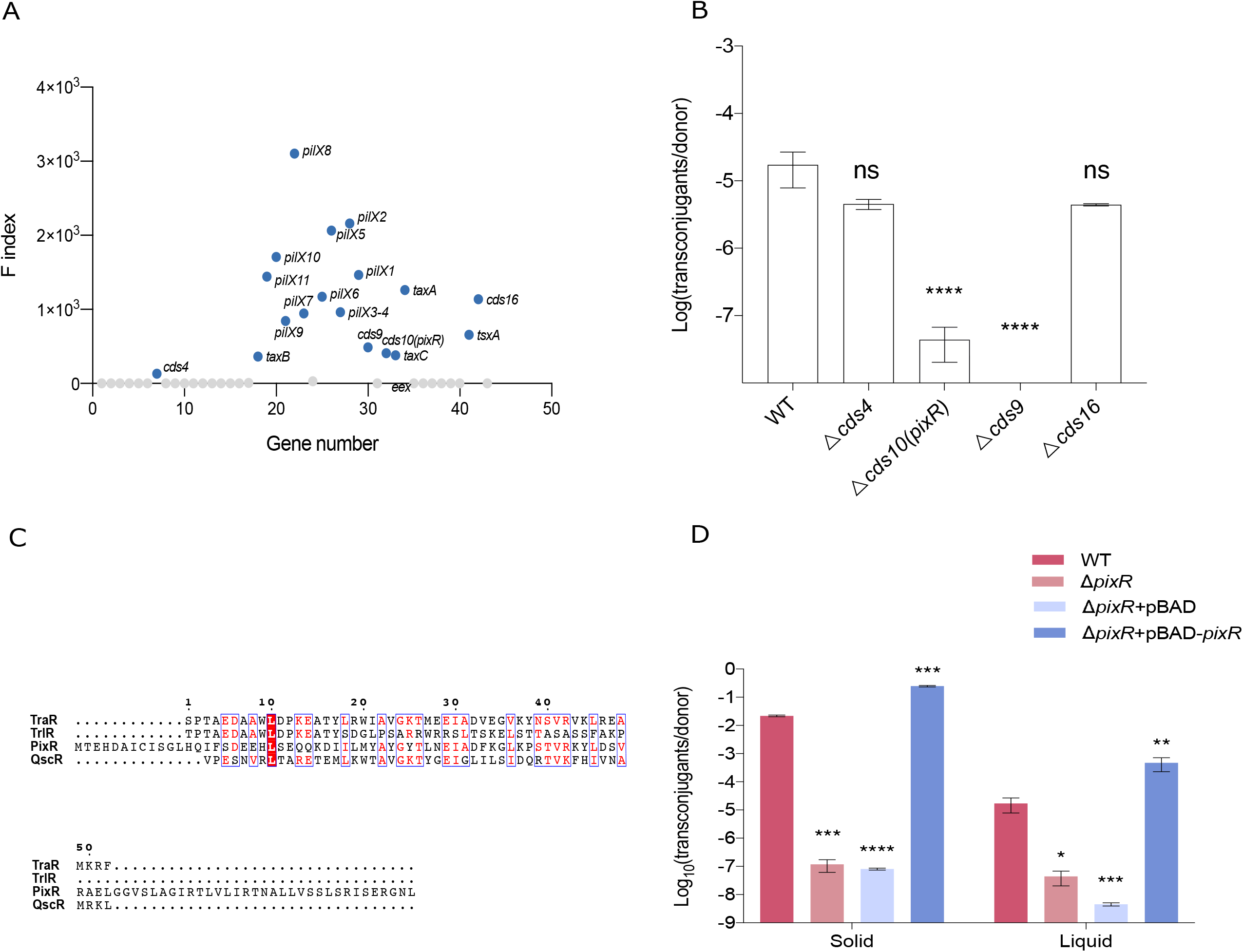
Identification of genes contributing to IncX4 conjugation. A) F index for every gene of pHNSHP23, based on HDTM data. F indexes were calculated as the insertion index ratio of the normalized transposon insertion read counts in the transconjugants mutant library divided by that in the initial mutant library. The genes with an F index > 100 were considered essential for conjugation and were highlighted in blue. The F indexes of *pilX*, *taxAC*, and *taxB* genes were used to set the threshold. B) Conjugation frequencies of pHNSHP23Δ*cds4*, pHNSHP23Δ*cds9*, pHNSHP23Δ*cds10*, pHNSHP23Δ*cds16*. The bars each represent the mean and standard deviation of biological triplicates. A one-way ANOVA with Tukey’s multiple comparison test was performed. C) Multiple sequence alignment of PixR with three members of the LuxR family of transcriptional regulators. TraR (Genbank WP_001278699), TrlR (Genbank CUX06573), and QscR (Genbank WP_088170053) are transcriptional regulators involved in quorum sensing from *A. tumefaciens*, *P. aeruginosa*. D) Effect of PixR on IncX4 plasmids conjugation under different conditions. Conjugation assays conducted on solid plates are shown on the left, whereas those performed overnight in LB broth are shown on the right. The bars each represent the mean and standard deviation of biological triplicates. Bars were compared using two distinct one-way ANOVA with Tukey’s multiple comparisons test. Statistical significance is indicated as follows: ****, P<0.0001; ***, P<0.001; **, P<0.01; *, P<0.05; ns, not significant.

### PixR is a key activator controlling IncX4 plasmid transfer

We then focused on the hypothetical gene *cds10* because its deletion strongly impaired the transfer of pHNSHP23 and it is a conserved feature of IncX4 plasmids, missing in other IncX subgroups. Hence, we hypothesized that *cds10* is an IncX4-specific conjugation factor. Based on the evidence presented below suggesting that *cds10* encodes a regulator of the expression of the *pilX* operon, this gene was renamed *pixR* (*pilX* regulator). The gene *pixR* is located directly downstream of *taxC* and upstream of the pilus assembly related genes (Figure 1). *pixR* encodes a 10.8-kDa hypothetical protein of 97 amino-acid residues with a predicted HTH DNA-binding domain (Pfam PF08281). The predicted structure of PixR obtained using phyre2 (http://www.sbg.bio.ic.ac.uk/phyre2/) suggests that PixR_3-79_ folds like the C-terminal domain of LuxR-type regulators despite low primary sequence identity (13~29%) (Figure 2C). LuxR proteins act as pheromone receptors and transcriptional regulators, affecting bacterial survival and propagation, virulence, and biofilm formation (31). The C-terminal domain of LuxR-type proteins typically binds specific DNA sites upon binding of an autoinducer (AI) to its N-terminal domain (31). However, PixR seems to lack the ligand-binding domain.

To further demonstrate the role of *pixR* in conjugation, we tested and compared the transfer efficiency of pHNSHP23 and its Δ*pixR* mutant in mating assays conducted both on agar plates and in broth. pHNSHP23 transferred at much higher rates on agar than in broth (2.19×10^-^ vs 1.73×10^-5^, respectively). pHNSHP23Δ*pixR* exhibited a 2-log reduction of transfer frequency in broth (1.73×10^-5^ vs 1.25×10^-7^) and a ~5-log reduction on agar (2.19×10^-2^ vs 1.18×10^-7^) (Figure 2D), suggesting that PixR acts as a key conjugation regulator both on a solid surface and in liquid. Transfer of the Δ*pixR* mutant was restored and even enhanced above the wild-type level upon overexpression of *pixR* from the arabinose-inducible promoter *P_BAD_* (Figure 2D). This observation indicates that *pixR* acts as a potent enhancer of the transmissibility of IncX4 plasmids.

### Overexpression of *pixR* up-regulates the transcription of the transfer genes

To explore how *pixR* regulates the transfer of IncX4 plasmids, we performed an RNA-seq experiment to determine the transcriptional profile of pHNSHP23 and its Δ*pixR* mutant with or without overexpression of *pixR* in *E. coli* BW25113. We found that the expression of 15 genes within a ~15-kb region increased considerably upon overexpression of *pixR* (Figure 1). Differential expression analyses confirmed that overexpression of *pixR* resulted in a 4.7 to 250-fold increase of expression of the *pilX* genes (Figure 3A). The expression of *taxB* and *trbM* also increased by 4.8-fold and the expression of three genes of unknown function (*cds6, cds8* and *cds9*) increased by 13-fold (Figure 3A). These observations suggest that PixR activates the transcription of a set of genes involved in the transfer of IncX4 plasmids. However, the expression levels were virtually identical between pHNSHP23 and its Δ*pixR* mutant (Figure 3B). Perhaps *pixR* is not expressed under the laboratory conditions of RNA extraction in pure culture without a recipient strain. Alternatively, the translation product of *pixR* is inactive or sequestered, preventing activation of transfer functions.

**Figure 3.**
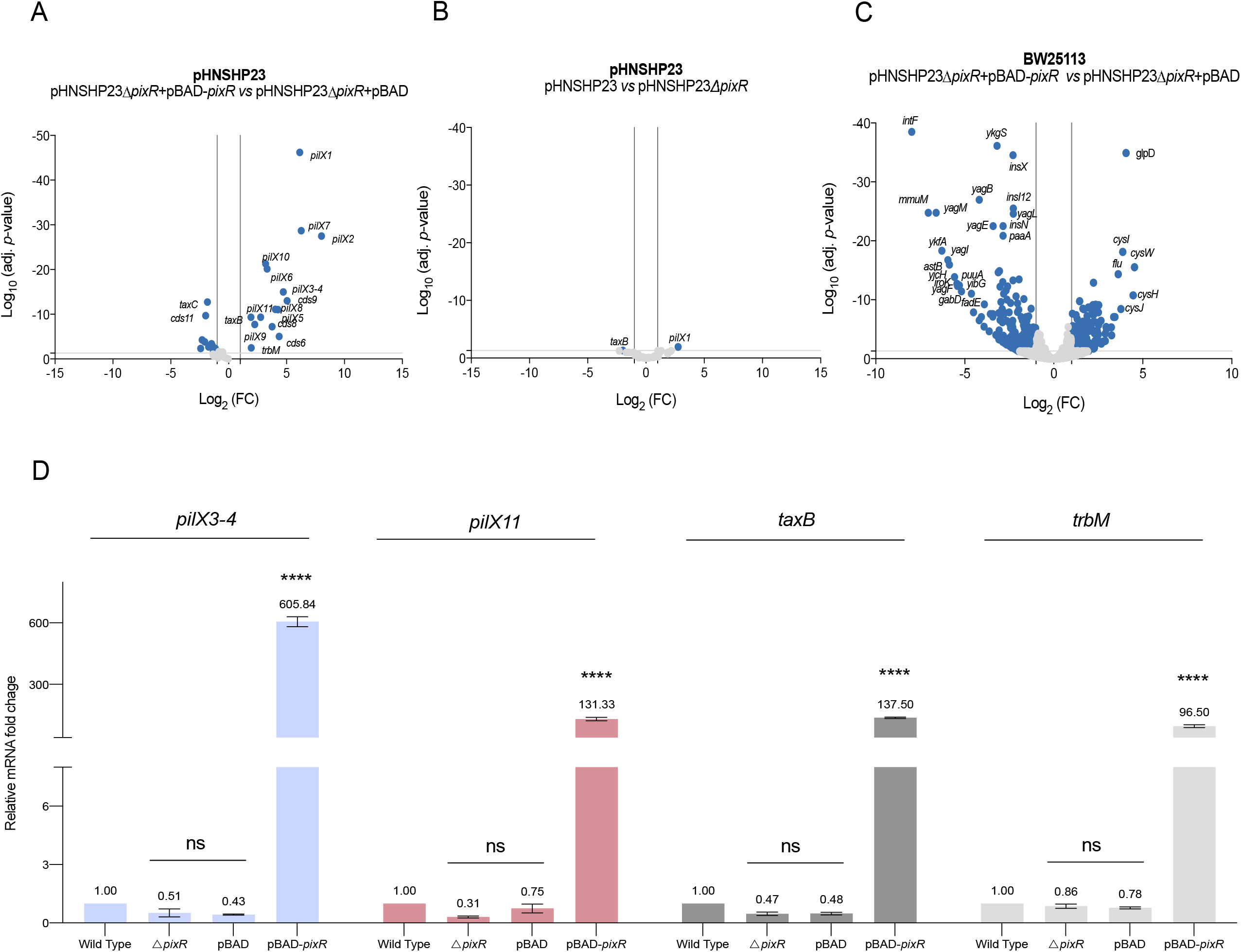
PixR regulon. A and B) Differential expression analyses of pHNSHP23 genes upon *pixR* overexpression or deletion. C) Differential expression analysis of BW25113 genes upon *pixR* overexpression. The X and Y axes represent the Log2 of fold change and the adjusted *p*-value, respectively. The vertical lines represent a 2-fold change in expression. Genes were considered as differentially expressed (and therefore marked as blue dots) when they displayed a fold-change of more than 2 as well as an adjusted *p*-value of more than 0.05, indicated by the gray horizontal line. D) Effect of *pixR* deletion on the mRNA level of *pilX3-4, pilX11, taxB*, and *trbM*. The bars each represent the mean and standard deviation of biological triplicates. Four distinct one-way ANOVAs with Tukey’s multiple comparisons test were performed to compare the relative mRNA levels of a given gene or operon under different conditions. Statistical significance is indicated as follows: ****, P<0.0001; ns, not significant.

Five hundred and eight chromosomal genes were differentially expressed upon *pixR* overexpression (Figure 3C). A pathway enrichment analysis showed that two pathways are up-regulated upon overexpression of *pixR*: ribosome (eco03010) and aminoacyl-tRNA biosynthesis (eco00970). Two pathways are also down-regulated: phenylalanine metabolism (eco00360) and flagellar assembly (eco02040) (Table S4). Surprisingly, several of the most down-regulated chromosomal genes belong to the cryptic prophage CP4-6 (Table S5).

We performed reverse-transcription followed by quantitative PCR (RT-qPCR) to confirm the RNA-seq data. Consistent with the results above, the mRNA levels of *pilX3-4, pilX11, taxB* and *trbM* increased more than 96-fold upon overexpression of *pixR* (Figure 3D). In contrast, a change in expression of less than 3-fold could be observed for the same set of genes when comparing pHNSHP23 with its Δ*pixR* mutant (Figure 3D).

### PixR directly activates the *pilX* operon

The presence of a predicted HTH DNA-binding domain in PixR suggests that PixR regulates the transcription of the transfer genes by binding to the promoter region upstream of *pilX1*. We used Cappable-seq to identify the transcriptional start sites (TSS) and compared the resulting profiles with or without *pixR* overexpression. A single peak located at 26,338 bp was observed exactly 42 bp upstream of *cds9* when *pixR* was overexpressed but not with the empty vector control, indicating that the RNA polymerase complex is recruited at this locus upon overexpression of *pixR*. No other peak was reliably detected downstream in the transfer region, suggesting that the genes *cds9, pilX1-6, cds8, pilX7-11, taxB* and *trbM* are all part of a single long operon, here referred to as the *pilX* operon (Figure 1). This organization is reminiscent of the *virB* genes of the archetypical P-type T4SS encoded by *Agrobacterium tumefaciens* which also constitutes a single operon (24, 32). However, the *pilX* operon of pHNSHP23 also includes *trbM* and *taxB* (Figure 4A). To further confirm this, a reverse transcription experiment was performed using a primer located at the 3’ end of *trbM* and total RNA extracted from *E. coli* BW25113 bearing pHNSHP23. A PCR amplification using the reverse cDNA confirmed that *cds9, trbM*, and *taxB* are, indeed, part of the *pilX* operon (Figure 4B and Table S3).

**Figure 4.**
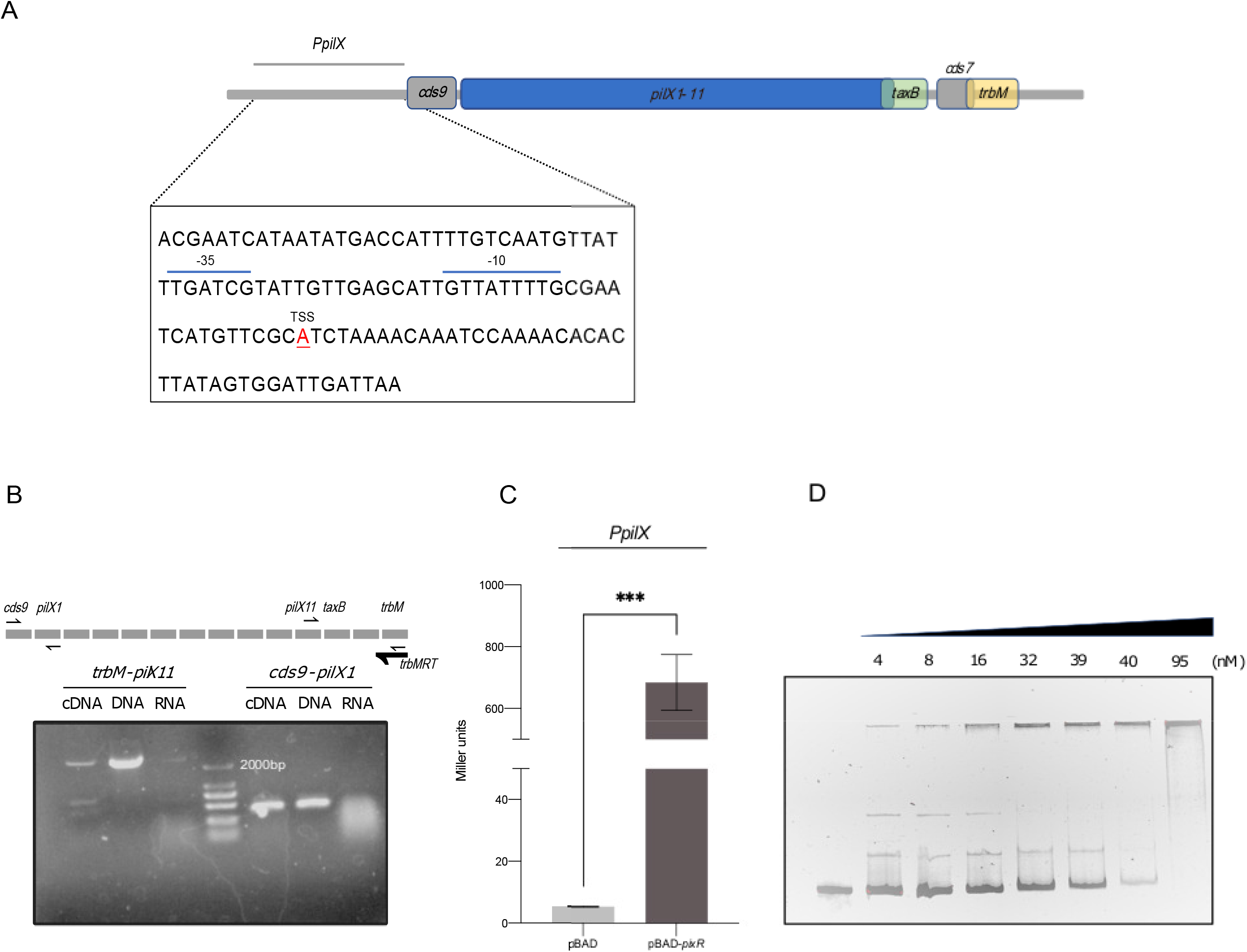
PixR directly activates the promoter of the *pilX* operon. A) Schematic representation of the promoter of the *pilX* operon. The transcription start site (TSS) was identified by Cappable-seq. The −10 and −35 boxes were predicted by Softberry (www.softberry.com). B) 1%agarose gel loaded with the amplified fragments of *cds9-pilX1* and *trbM-pill11*. A primer located at the 3’ of *trbM* was used for reverse transcription. gDNA and RNA were used as positive and negative controls, respectively. C) β-galactosidase activities of *P_pilX_*. The activity of *P_pilX_* was monitored from a transcriptional *lacZ* fusion (*P_pilX_-lacZ*) in *E. coli*. Cultures were grown in LB broth with 0.2% arabinose at 37 °C for 2 hours to induce the expression of *pixR*. The bars each represent the mean and standard deviation of biological triplicates. They were compared using an unpaired *t*-test (***, P<0.001). D) EMSA performed with the *P_pilX_* fragment and PixR. The 6×His-tagged PixR protein was purified by Ni-NTA affinity chromatography. Increasing concentrations of PixR protein were incubated with the *P_pilX_* fragment for 1 hour.

The 120-bp region upstream of the ATG start codon of *cds9*, named *P_pilX_*, was cloned upstream a promoterless *lacZ* gene (Table S3). The β-galactosidase activity was measured with and without ectopic expression of *pixR* in *E. coli* BW25113 in the absence of pHNSHP23. Without *pixR*, we detected a weak β-galactosidase activity for *P_pilX_* that increased 130-fold upon *pixR* overexpression (Figure 4C), confirming that PixR directly activates *P_pilX_*.

To test whether PixR directly binds onto *P_pilX_*, we performed an Electrophoretic Mobility Shift Assay (EMSA) using the purified PixR protein. The 120-bp PCR-amplified *P_pilX_* probe was incubated with an increasing concentration of purified PixR protein. The observed size shift of the fragment confirmed that PixR binds to *P_pilX_* (Figure 4D).

Altogether, these results demonstrate that PixR binds to the promoter region of the *pilX* operon and, by doing so, directly activates its transcription with very high efficiency *in vitro*.

### The ecological role of *pixR* in pHNSHP23

Plasmid persistence in a bacterial community is usually associated with efficient acquisition and fitness benefits. To evaluate the ecological role of *pixR* in the persistence of pHNSHP23, we compared the ability of pHNSHP23 or its Δ*pixR::cat* (Cm^R^) mutant to invade a plasmid-free population. The Δ*pixR* plasmid failed to invade the plasmid-free population and was progressively lost after the first day (Figure 5A). In contrast, pHNSHP23 gradually invaded the plasmid-free cells until the concentration of pHNSHP23-harboring cells equaled that of plasmid-free cells (Figure 5B). The population dynamics in competition co-cultures were consistent with the observation in individual plasmid invasion assays (Figure 5C). Worthy of note, cell growth kinetics and pHNSHP23 stability remained unaffected by the Δ*pixR* mutation over 10 days in pure culture (Figures S1B and S1C). For this reason, and because the Δ*pixR* plasmids are lost and fail to invade a population of plasmid-free cells, we hypothesized that the deletion of *pixR* induces a fitness cost. Hence, we evaluated the competition capability of pHNSHP23 and its Δ*pixR* mutant. While pHNSHP23 conferred a fitness advantage to its *E. coli* host, the Δ*pixR* plasmid inflicted a fitness cost. Specifically, the average relative fitness decreased from 0.95 to 0.68 over the course of 5 days (Figure 5Da and Figure 5Db).

**Figure 5.**
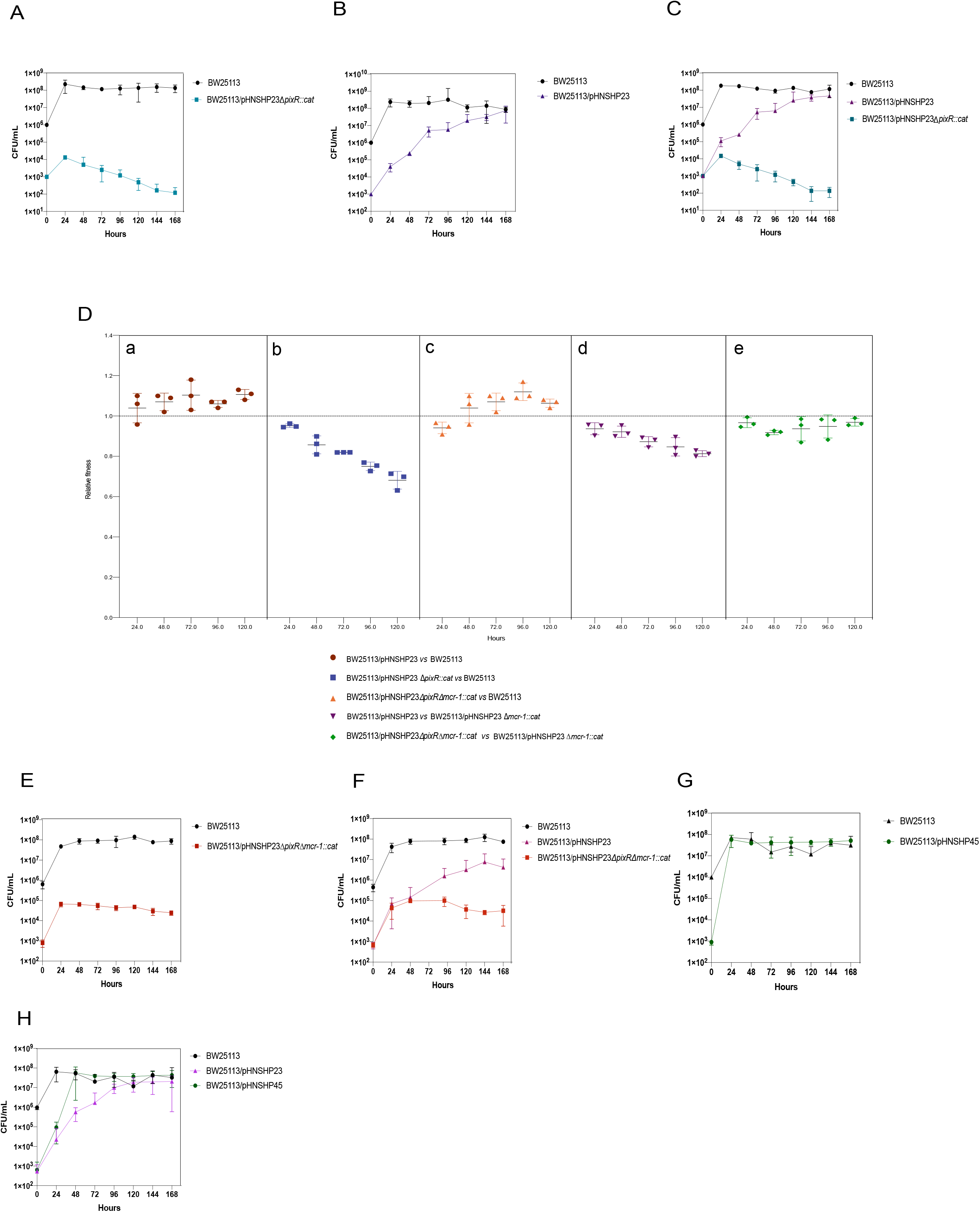
The ecological role of *pixR* for plasmid pHNSHP23 invasion and persistence. *E. coli* BW25113/pHNSHP23Δ*pixR::cat* (A), BW25113/pHNSHP23 (B), BW25113/pHNSHP23Δ*pixR*Δ*mcr-1::cat* (E) and BW25113/pHNSHP45 (G) were mixed with 1 000-fold excess of BW25113 without plasmid initially; co-cultures of BW25113/pHNSHP23Δ*pixR::cat* (C), BW25113/pHNSHP23Δ*pixR*Δ*mcr-1::cat* (F) or BW25113/pHNSHP45 (H) with BW25113/pHNSHP23 and 1 000-fold excess of BW25113 without plasmid initially. D) *E. coli* BW25113/pHNHSP23, *E. coli* BW25113/pHNSHP23Δ*pixR::cat* or BW25113/pHNSHP23Δ*pixR*Δ*mcr-1::cat* were competed with the reference strain BW25113 *in vitro*, separately. BW25113/pHNSHP23Δ*mcr-1* were competed with BW25113/pHNSHP23 *in vitro*. BW25113/pHNSHP23Δ*pixR*Δ*mcr-1*::*cat* was competed with BW25113/pHNSHP23Δ*mcr-1*::*cat in vitro*. All competitions assays were performed with three biological replicates.

We suspected that the fitness cost of pHNSHP23Δ*pixR* was associated with *mcr-1* carriage. Indeed, our previous results showed that even a single copy of *mcr-1* exerts a fitness burden upon *E. coli* (7). To test this hypothesis, we carried out a competition assay between pHNSHP23 and its Δ*mcr-1::cat* (Cm^R^) mutant. Compared to the Δ*mcr-1* mutant, pHNSHP23 showed a reduced relative fitness, decreasing from 0.93 to 0.81 over 5 days (Figure 5Dd). Together, these results confirm *mcr-1* carried by an IncX4 plasmid imposes a fitness cost to *E. coli*.

We then tested the invasion and competition capability of pHNSHP23Δ*pixR*Δ*mcr-1::cat.* As expected, the Δ*pixR* Δ*mcr-1* mutant failed to invade a population of plasmid-free cells, but the population harboring the plasmid remained stable (Figures 5E and 5F). Additionally, the Δ*pixR* Δ*mcr-1* mutant conferred a slight fitness advantage to *E. coli*, though its fitness was not as high as that of the Δ*mcr-1* mutant (Figure 5Dc and Figure 5De). These results suggest that *pixR* compensates for the fitness burden inflicted by *mcr-1*.

Collectively, these results suggest that the increased rate of conjugation aids IncX4 plasmids in invading and persisting in bacterial populations. Because IncX4 and IncI2 plasmids are the main drivers of *mcr-1* spread, we compared their invasion capabilities using pHNSHP23 and pHNSHP45 as models, respectively. We found that pHNSHP45 invades cells at a much higher rate than pHNSHP23. pHNSHP45 had invaded most cells after 24 h in individual invasion conditions and after 48 h in competition cultures (Figures 5G and 5H). In contrast, pHNSHP23 failed to overtake plasmid-free cells even after 5 days (Figures 5B and 5H). The higher transfer rate reported for IncI2 plasmids (~10^-1^-10^-2^) compared to IncX4 plasmids (~10^-2^-10^-4^) could explain these results (4).

### Comparative genomics of IncX plasmids

We then searched the GenBank database for PixR protein homologues using blastp and found that 219 out of 247 IncX4 plasmids encode PixR (Table S1), while the other 28 IncX4 plasmids encode two types of PixR-like proteins (PixR-1 and PixR-2) sharing 46% identity with PixR of pHNSHP23 (Figure S1). Beyond the IncX4 family, a putative regulatory protein encoded by IncX7 plasmids p3 and pJARS35 from *Yersinia pestis* shares 54% identity with PixR. In addition, two PixR-like proteins encoded by two Col plasmids and other small untyped plasmids also share 44% and 49% identity with PixR. No other PixR-like proteins were identified in other IncX plasmids (Figure S1).

Considering *pixR* is a transfer activator specific to IncX4 and IncX7 plasmids, we compared the transfer genes of different IncX subgroups. We found that the conjugative transfer locus *taxB-pilX-taxCA* of the archetypical IncX2 plasmid R6K (Genbank AJ006342) shares >68% identity with the corresponding locus of the other representative IncX plasmids pOLA52 (IncX1a, EU370913), R485 (IncX1b, HE577112), pEC14_35 (IncX3, JN935899), pBK31567 (IncX5, JX193302), and pCAV1043-58 (IncX8, CP011588), except for the entry exclusion gene *eex* of IncX1b plasmid R485. In contrast, the corresponding locus of plasmids pHNSHP23 (IncX4) and p3 (IncX7, CP009993), while exhibiting >70% identity with each other, share less than 25% identity with plasmids of the other IncX groups (Figure 6). Furthermore, *eex*, usually found between *pilX5* and *pilX6* in most IncX subgroups, is located upstream of *pilX1* in pHNSHP23 and p3. Finally, *pixR* in pHNSHP23 and p3 is missing in other IncX subgroup plasmids where *actX* is found instead (Figure 6 and Table S6). Pairwise sequence comparison showed no significant similarities between *actX* and *pixR*.

**Figure 6.**
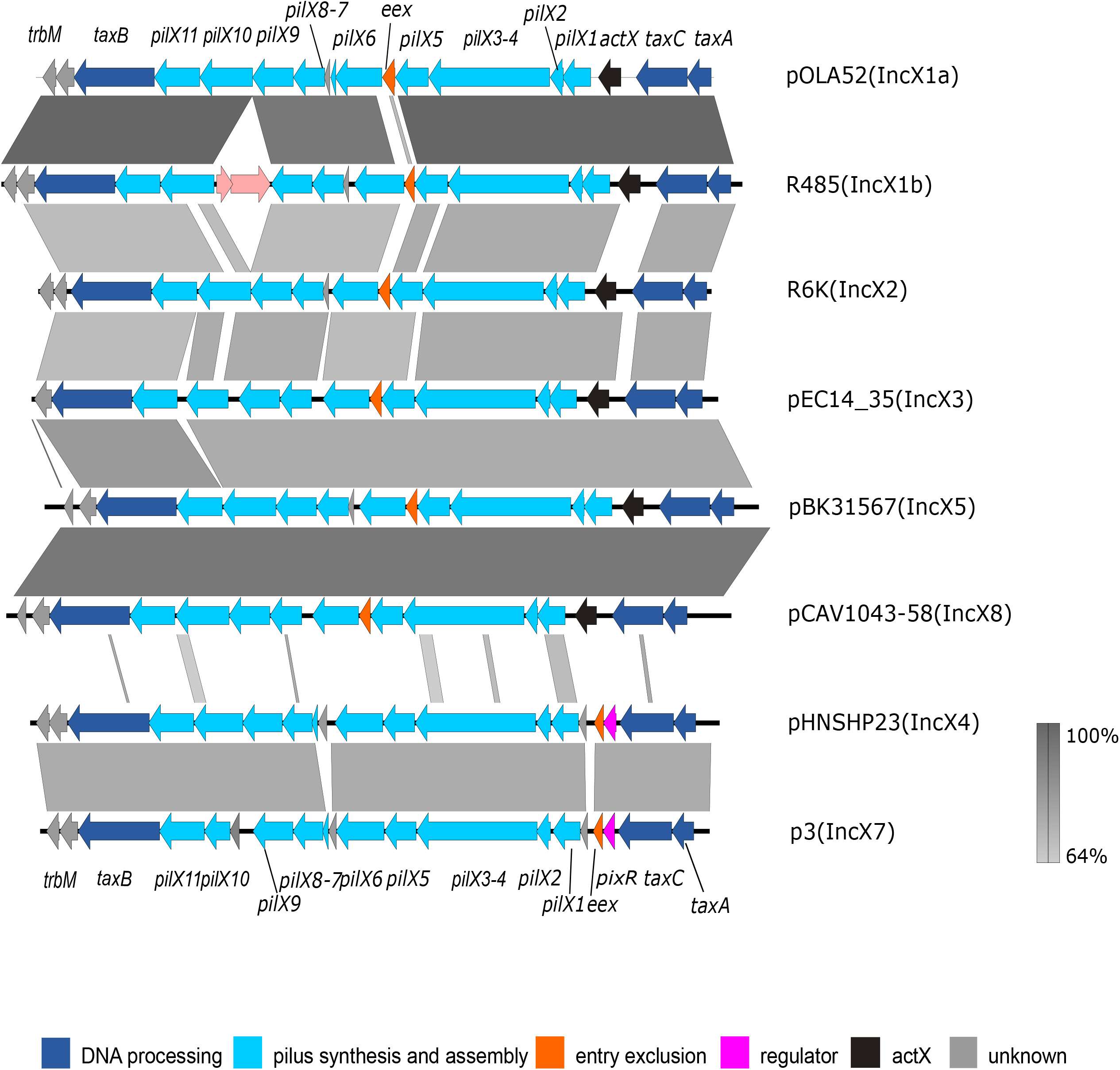
Linear comparison of IncX plasmids transfer region. Genes are labeled in different colors based on their annotation; Shades of gray indicate the percentage of sequence identity.

## DISCUSSION

IncX4 plasmids carrying *mcr-1* have been detected in over 41 countries and regions. Given the central role of conjugative plasmids in the global spread of antibiotic resistance, understanding the factors contributing to the dissemination of *mcr-1*-bearing IncX4 plasmids is imperative. In this study, we screened systematically for genes involved in the conjugative transfer of IncX4 plasmids and have identified the novel transfer activator PixR that contains the conserved C-terminal DNA binding domain of LuxR proteins. We showed that PixR directly activates the expression of a set of core transfer genes by binding to the promoter of the transfer operon. Using HDTM, we identified 18 genes in pHNHSP23 that are essential for conjugative transfer. Thirteen of them were found to be up-regulated upon overexpression of *pixR*, indicating that *pixR* plays an important role in the regulation of IncX4 conjugation. To the best of our knowledge, few studies have explored the regulation of the conjugative transfer of IncX plasmids. In IncX3 plasmids, the absence of a plasmid-encoded H-NS-like protein up-regulates the expression of transfer genes through an uncharacterized pathway, causing a 2.5-fold increase in transfer frequency of IncX3 plasmids (33). Thus, PixR is the first transcriptional activator of transfer genes identified in IncX plasmids, which expands our knowledge of the conjugation of IncX plasmids. In the Ti plasmid of *A. tumefaciens*, the PixR homologue TraR also belongs to the LuxR family of proteins and up-regulates the expression of the T4SS-encoding *traI-trbCDEJKLFGHI* operon, leading to the activation of transfer (34, 35). However, in the case of TraR, binding of the 3-oxo-octanoyl-homoserine lactone (OC8HSL) to the N-terminal domain of TraR is required for activation (36). In contrast, our data seem to indicate PixR lacks the ligand-binding domain. Further studies are needed to explore its activation mechanism. In particular, the signals that activate the expression of *pixR* are still unknown. The sequence divergence of IncX4 and IncX7 plasmid transfer regions with those of other IncX subgroups and the substitution of *actX* for *pixR* in other subgroups suggest that the PixR-based regulatory mechanism is specific to IncX4 and IncX7 plasmids. The genetic determinants that regulate the conjugal transfer of other IncX subgroups still need to be identified. The presence of *pixR* in Col or small untyped plasmids suggests a common regulatory mechanism in these plasmids and the possibility of co-regulation if these plasmids co-exist in the same host.

The conservation of *pixR* in the IncX4 plasmids suggests that this gene provides an evolutionary advantage for *mcr-1*-bearing IncX4 in nature. *pixR* increases the transfer capability of IncX4 plasmids and a high rate of conjugation usually helps plasmids invade bacterial populations (19, 20, 37). Therefore, here we investigated the role of *pixR* in the persistence and invasion of *mcr-1*-bearing IncX4 plasmids to better understand their prevalence. We found that pHNSHP23 lacking *pixR* inflicts a fitness cost to an *E. coli* host after 24 h of culture. The inactivation of *mcr-1* lessened this burden. These results demonstrate that efficient conjugation is sufficient to overcome the fitness cost of low-copy *mcr-1* carriage. The presence of *pixR* increases the transfer capability of IncX4 plasmids, minimalizing the fitness cost exerted by *mcr-1* carriage, resulting in the successful dissemination of *mcr-1*-bearing IncX4 plasmids. Through plasmids invasion assays, we found that pHNSHP23 lacking *pixR* loses its invasion capability and fails to establish stably in the cell population, suggesting that *pixR* is important for the survival of *mcr-1*-bearing IncX4 in a bacterial community (Figure 7). Even though pHNSHP23 was able to invade a plasmid-free population, the rate of invasion was much slower than that of IncI2 plasmid pHNSHP45, which overtakes a plasmid-free population within 24-48 hours. Such difference is likely caused by a higher rate of transfer of IncI2 plasmids compared to IncX4 plasmids (4). This could also explain the lower detection rate of IncX4 plasmids compared to IncI2 plasmids after the withdrawal of colistin used as a growth promoter in China (5). Overall, our data highlight that the efficient conjugation of plasmids plays an important role in the prevalence and persistence of *mcr-1*-bearing plasmids. The strict controlling of low plasmid copy number as well as efficient conjugation both contribute to the global prevalence of *mcr-1*-bearing plasmids (7).

**Figure 7.**
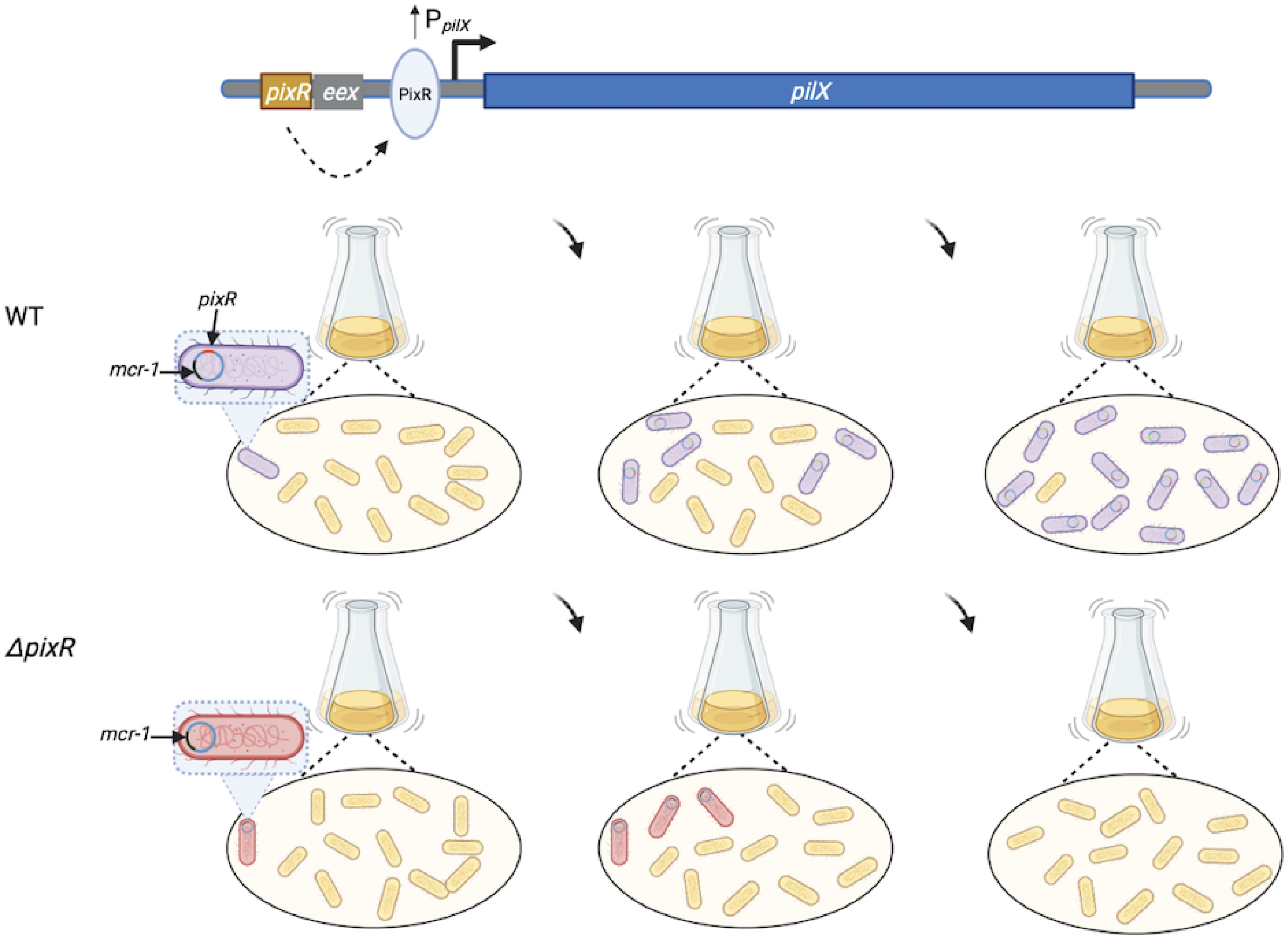
PixR’s key role in the persistence and invasion of *mcr-1*-bearing IncX4 plasmids. A) PixR enhances the conjugation efficiency of IncX4 plasmids by binding the promoter of a set of essential transfer genes to increase their expression. B) PixR has a strong effect on the persistence of *mcr-1*-bearing IncX4 plasmids in bacterial populations.

Curiously, a limited (1~3-fold) difference of *pilX3-4* and *pilX11* expression was observed between wild-type pHNSHP23 and its Δ*pixR* mutant in RNA-seq and RT-qPCR data, suggesting that PixR is not active or sequestered in non-mating conditions. Alternatively, *pixR* expression is possibly repressed by another unknown factor in *E. coli* under laboratory conditions. This result is consistent with previous reports showing that repression of the conjugative transfer functions enhances the fitness of the bacterial host. Transfer genes are usually repressed with only a few cells in the population expressing the conjugative machinery (34, 38, 39). A plausible hypothesis could be the transient elicitation of *pixR* expression or PixR activity in a small subset of cells in the donor population upon contact with a potential recipient cell, such as a former donor cell that has lost pHNSHP23 due to *mcr-1* carriage. *pixR* would then compensate the cost associated with *mcr-1* by enhancing conjugative transfer for a limited number the donor cells to maintain the balance of plasmid gain/loss. As mentioned above, future studies will need to explore the activation mechanism of PixR.

In conclusion, we characterized an IncX4-specific regulatory mechanism that controls plasmid conjugation and showed that efficient conjugation is important to alleviate the fitness cost of *mcr-1* carriage and promote the persistence and invasion of *mcr-1*-bearing plasmids within a bacterial population. Given the significant role of high conjugation frequency of plasmids in the spread of antibiotic resistance genes, plasmid transfer inhibitors are noble approaches for solving the antibiotic resistance crisis. A deeper knowledge of the regulation of plasmid conjugation will provide potential targets for conjugation inhibition. Our data provide a new target to inhibit the dissemination of antibiotic resistance genes, especially *mcr-1*, mediated by IncX4 plasmids.

## SUPPLEMENTARY DATA

Supplementary Data are available Online

## ACKNOWLEDGMENTS

This work was supported by the National Natural Science Foundation of China (grants no.31830099 and 31625026), Guangdong Special Support Program Innovation Team (No. 2019BT02N054), the 111 Project (No. D20008), and the Innovation Team Project of Guangdong University (No. 2019KCXTD001).

Discovery grant [RGPIN-2016-04365] from the Natural Sciences and Engineering Research Council of Canada (NSERC) and Project grant [PJT-153071] from the Canadian Institutes of Health Research (CIHR) to V.B. R.D. is the recipient of a Fonds de recherche du Québec-Nature et Technologies (FRQNT) doctoral fellowship.

## AUTHOR CONTRIBUTIONS

J-HL, VB and LXY designed the study. LXY, JY and KYY performed experiments. LXY, RD, GF, VB and J-HL analyzed the data and prepared the figures. LXY, J-HL, VB, RD, and GF drafted the manuscript. All authors reviewed, revised, and approved the final manuscript.

## LEGENDS FOR SUPPLEMENTARY MATERIAL

**Text S1.** Additional experimental procedures. (DOCX)

**Figure S1**. a. Geographic distribution of IncX4 plasmids. Countries where IncX4 plasmids were more prevalent are colored with a darker shade of blue. b. Growth curves of E. coli BW25113 carrying pHNSHP23 and pHNSHP23ΔpixR mutants. c. In vitro stability of pHNSHP23 and pHNSHP23ΔpixR in E. coli BW25113. d. Protein sequence alignment of PixR and PixR-like proteins. The amino acid sequences of PixR and PixR-like proteins encoded by the representative plasmids were aligned using MUSCLE 3.8.31. PixR1 and PixR2 correspond to homologues encoded by 28 IncX4 plasmids. (DOCX)

**Table S1.** Characteristics of IncX4 plasmids in Genbank database. (XLSX)

**Table S2.** Strains and plasmids used in this study. (DOCX)

**Table S3.** Oligonucleotides used in this study. (DOCX)

**Table S4.** Up- and down-regulated BW25113 KEGG pathways. (DOCX)

**Table S5.** RNAseq transcriptome profiling of BW25113. (XLSX)

**Table S6.** Comparison of pHNSHP23 conjugative transfer proteins with their respective homologues encoded by other IncX representative plasmids. (DOCX)

